# Continuous extraction and concentration of secreted metabolites from engineered microbes using membrane technology

**DOI:** 10.1101/2022.03.10.483787

**Authors:** Sebastian Overmans, Gergo Ignacz, Aron K. Beke, Jiajie Xu, Pascal Saikaly, Gyorgy Szekely, Kyle J. Lauersen

## Abstract

Microalgal cultivation in photobioreactors and membrane separations are both considered sustainable processes. Here we explore their synergistic combination to extract and concentrate a heterologous sesquiterpenoid produced by engineered green algal cells. A hydrophobic hollow-fiber membrane contactor was used to allow interaction of culture broth and cells with a dodecane solvent phase to accumulate algal produced patchoulol. Subsequent continuous membrane extraction of patchoulol from dodecane enabled product concentration in a methanol stream as well as dodecane recovery for its reuse. A structure-based prediction using machine learning was used to model a process whereby 100% patchoulol recovery from dodecane could be achieved with solvent-resistant nanofiltration membranes. Solvent consumption, E-factor, and economic sustainability were assessed and compared with existing patchoulol production processes. Our extraction and product purification process offers six- and two-orders of magnitude lower solvent consumption compared to synthetic production and thermal-based separation, respectively. Our proposed methodology is transferable to other microbial systems for the isolation of high-value isoprenoid and hydrocarbon products.

## Introduction

The strive to create environmentally sustainable processes in the chemical and biochemical industries has led to the definition of the twelve principles of green chemistry, which has also been extended to chemical engineering and bio-processes.^1^ Sustainable sourcing of chemicals is important for global drives towards resource circularity and a reduction of the environmental impact of human activities. Transfer of genetic modules for specialty metabolism from a progenitor organism into an easier to handle microbial system through synthetic biology now allows the production of otherwise difficult to scale specialty chemicals in controlled bio-processes^2^. One of the most established fields in synthetic biology is metabolic engineering of heterologous isoprenoid production^3, 4^. Fermentative bacterial or yeast cell expression systems have been commonly used for this purpose, owing to their genetic tractability. Recently, the green alga *Chlamydomonas reinhardtii* has also emerged as an alternative and more sustainable isoprenoid production host, with demonstrated sesqui- and diterpene production^5-8^. Some microalgae, like *Botryococcus braunii*, also naturally secrete hydrocarbon compounds, including alkadienes, alkatrienes, or polyterpenoid botryococcenes^9^. Isoprenoid products are largely hydrophobic molecules and accumulate in small amounts in microbial hosts unless a physical sink is provided to generate pull and enable forward metabolic reactions^5^. This physical sink is achieved at lab scale with hydrophobic bio-compatible solvents like dodecane (C12)^10^. However, the microbial culture-solvent system is difficult to scale as hydrophobic parts of the microbial culture can form turbid emulsions with the solvent, limiting practical implementation^11^. In order to scale such systems to sustainable bio-processes, stable culture-solvent interactions must be developed and coupled to efficient downstream processes which minimize chemical waste generation in the isolation of target chemicals.

Hollow-fiber (HF) nanofiltration membranes enable high-surface-area interactions of aqueous and solvent phases in compact modules. Here, we investigated the use of hydrophobic HF membrane contactors to act as a high-surface-area, physical interaction matrix between dodecane solvent and engineered green algal cells to continuously extract the heterologous isoprenoid product patchoulol. Upon extraction, patchoulol accumulates in the dodecane solvent, which requires further processing to yield the target product. Further concentration and purification must be efficient and generate as little waste as possible. We then further investigated coupling our algal isoprenoid extraction process to organic solvent nanofiltration (OSN) as an efficient downstream process module to recover dodecane solvent and isolate the patchoulol product. OSN is a continuous, pressure-driven, low energy, low waste, liquid-liquid, or liquid-solute membrane separation technology.^12, 13^ Chemical concentration, catalyst and solute recovery, solvent recycling, and product purification have been accomplished using OSN in the petrochemical and pharmaceutical industries. Natural product extraction^14-16^ and concentration^17^, as well as biomass processing^18^, are emerging applications of OSN.

Continuous liquid–liquid separators have been used to connect microflow reactions with separation modules^19^. These separators are based on the different wettability properties of the solvents on membranes. A typical liquid–liquid separator (for example a Zaiput module) contains an ultrafiltration membrane with strong hydrophilic or hydrophobic properties^20^. These ultrafiltration membranes are chosen to be wetted by one of the solvents from the two-phased system but not by the other. This selective wetting property allows one solvent to permeate through the membrane while the other is retained. Careful selection of the solvents, flowrate and the membranes are crucial for a successful separation. They use less energy compared to conventional thermal-based separation methods, such as distillation or crystallization; or chromatography-based separations. The implementation of a continuous separation module also allows recovery of the solvents to be recycled within the process without the need of additional process steps. Continuous liquid–liquid separation can be considered a low-cost, low-energy alternative to conventional separation techniques^21^.

Design for separation engineering as a field develops bio- and chemical processes so that product separation feasibility and overall efficiency are considered in every design decision. The work presented here employs process design elements, machine learning, and modeling tools to demonstrate a sustainable continuous isoprenoid extraction process from engineered green microalgal culture. The process could equally be applied to any engineered or natural microbe secreting hydrophobic products of chemical value. In this study, we demonstrate HF technology can be used for heterologous isoprenoid extraction, demonstrate a product concentration method with minimal waste generation, and model strategies to increase downstream bio-process efficiency.

## Experimental

### Materials

Ethanol (96% vol.), methanol, and n-dodecane (≥99%) were purchased from VWR International (Fontenay-sous-Bois, France). Prior to its use, dodecane was passed through a Supelclean^™^ LC-Si solid-phase extraction (SPE) column (Product no. 505374; Sigma-Aldrich, Taufkirchen, Germany) to remove impurities.

Microalgal cultures were routinely maintained in Tris acetate phosphate (TAP) medium^22^ with updated trace element solution^23^ using nitrate as a nitrogen source instead of ammonium. A microfiltration hollow-fiber (HF) cartridge with 750 hydrophobic polysulfone membrane fibers (surface area: 0.93 m^2^, pore size: 0.65 μm, product code CFP-6-D-9A; Cytiva, USA) was used to extract patchoulol from *C. reinhardtii* culture. IL-400 membranes for the SEP-10 membrane contactor were purchased from Zaiput (USA). Puramem S260 and Duramem 300 membranes for the nanofiltration were purchased from Evonik (Germany). The GMT-oNF-2 nanofiltration membrane was purchased from Borsig (Germany).

### Microalgal culture and hollow-fiber setup

To investigate the potential of hollow-fiber technology to extract terpenoid products from microbial cultures, the UVM4 strain^24^ of the green model microalga *Chlamydomonas reinhardtii* was engineered through previously reported genetic strategies^5^ to produce the sesquiterpenoid patchoulol and use nitrate as a sole nitrogen source.^25^ Before being used in the hollow-fiber, the culture was routinely maintained on TAP NO^3^ agar plates and then transferred into 24-well plates and subsequently into a 125 mL Erlenmeyer flask under growth conditions previously described^26^. 20 mL of dense culture was added to 1.98 L of TAP NO_3_ media in a 2 L Erlenmeyer flask. The flask was then connected to the HF cartridge to extract patchoulol from the algae (see Fig. 1 and Suppl. Fig. 1). The algal culture in the reservoir was stirred at 200 rpm and pumped through the shell side of the cartridge with a peristaltic pump (Masterflex L/S Series; Masterflex, USA) at a flow rate of 45 mL min^−1^. The solvent reservoir contained 400 mL of dodecane that was pumped through the lumen side in cross flow to the algae culture at a rate of 7.5 mL min^−1^ with a high-pressure pump (Azura P 4.1S; Knauer, Germany). Cells were allowed to accumulate on the HF membrane and were not washed off during medium exchanges, which were performed in the culture reservoir. The algae culture in the cartridge was illuminated with two standard white-light OLED panels (approx. 250 μmol m^−2^ s^−1^), and the HF bioreactor was maintained under those conditions for 60 d. Dodecane was sampled (500 μL) daily from the solvent reservoir for patchoulol quantification using gas chromatography-mass spectrometry (GC-MS) as described below.

**Figure 1.**
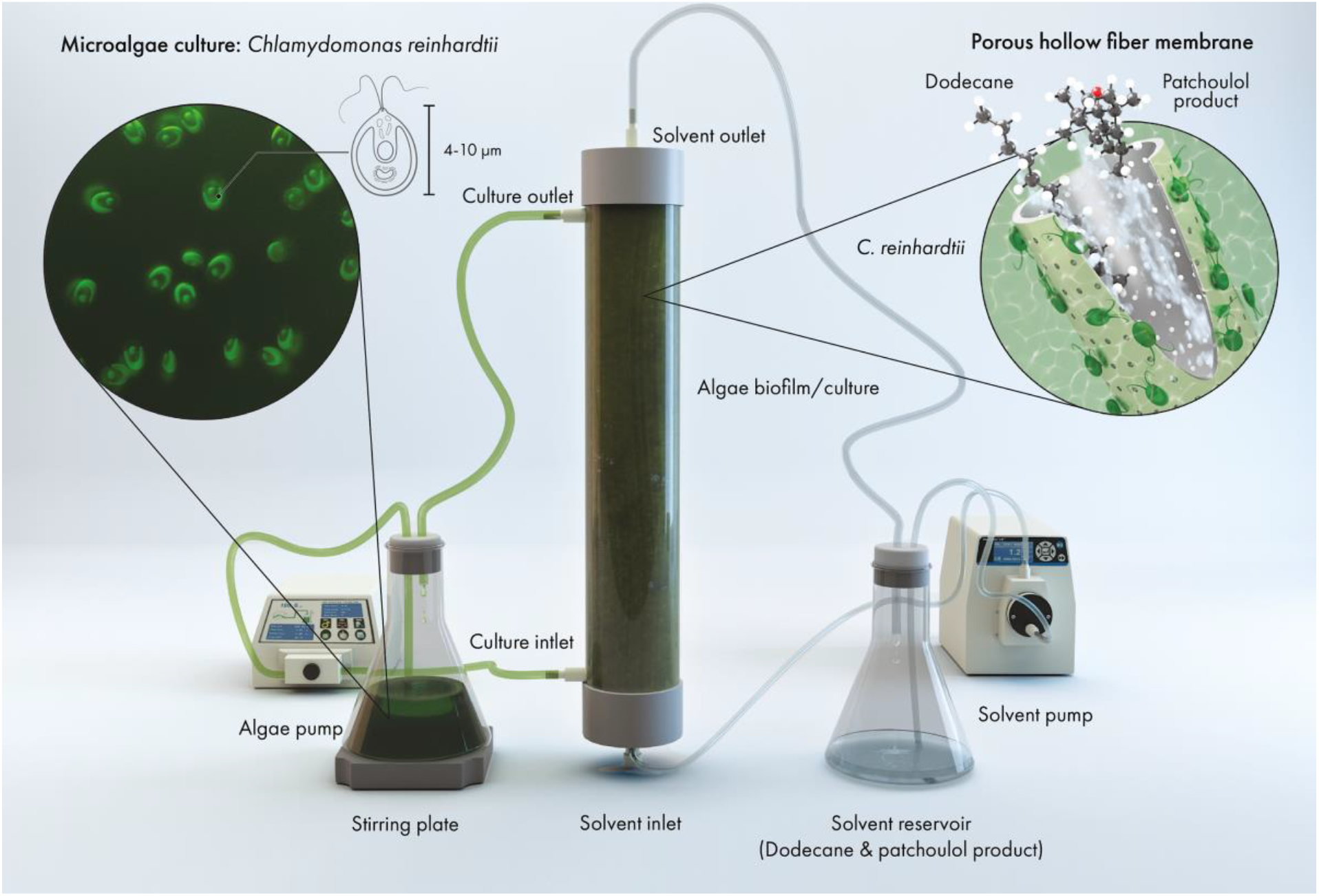
Illustration of the hollow-fiber setup used for the extraction of patchoulol from engineered *C. reinhardtii* culture using dodecane as solvent. Once the algae culture had established on the outside of the hollow fibers (shell side), dodecane was pumped in cross-flow to the culture through the hollow-fibers (lumen side). At the pores, dodecane comes in contact with the attached algae cells and medium, which initiates the extraction process of patchoulol. Figure produced by Ana Bigio, scientific illustrator.

### Chlorophyll fluorescence measurements

To determine the health of the algal culture inside the reservoir and the HF cartridge, variable chlorophyll fluorescence of photosystem II (PSII) was measured with a Pulse Amplitude Modulation (PAM) fluorometer (Mini-PAM-II; Heinz Walz GmbH, Germany). Before each measurement, the light source of the HF setup was switched off for 15 min to dark-adapt algae cells and to ensure open PSII reaction centers. The signal amplitude was adjusted before five single-turnover measurements (n=5) each were recorded of the algae inside the flask and at different locations around the HF cartridge. The maximum photochemical efficiency (*F*_v_/*F*_m_) was determined using Eqn 1, where *F*_m_ and *F*_0_ are the maximal and minimal PSII fluorescence of dark-acclimated *C. reinhardtii* cells, respectively ^27^:

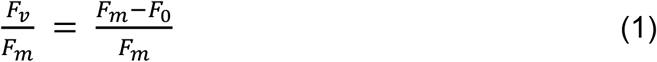

### Nutrient analysis

To monitor the concentrations of macronutrients in the culture medium, samples of the algae culture were taken daily. Specifically, 25 mL of culture was sampled daily from the reservoir flask. The sampled culture was centrifuged at 4000xg for 10 min (18 °C), and the clarified supernatant was filtered with a 0.2 μm syringe filter to remove insoluble particles. The filtrate was used for the determination of nitrate (Hach kit LCK339) and total phosphorous (TNT844) concentrations (in mg/L), following manufacturer’s protocols (Hach, Germany). Sample absorption was measured using a DR 1900 spectrophotometer (Hach, Germany). Each nutrient concentration was analyzed in biological duplicates (n=2). Where necessary, samples were diluted with fresh culture medium to be within linear analysis range.

The culture medium in the HF setup was regularly either replenished (100 mL added) or 95% of the liquid culture (1.9 L) was exchanged with fresh TAP medium (see Fig. 2c for schedule).

**Figure 2.**
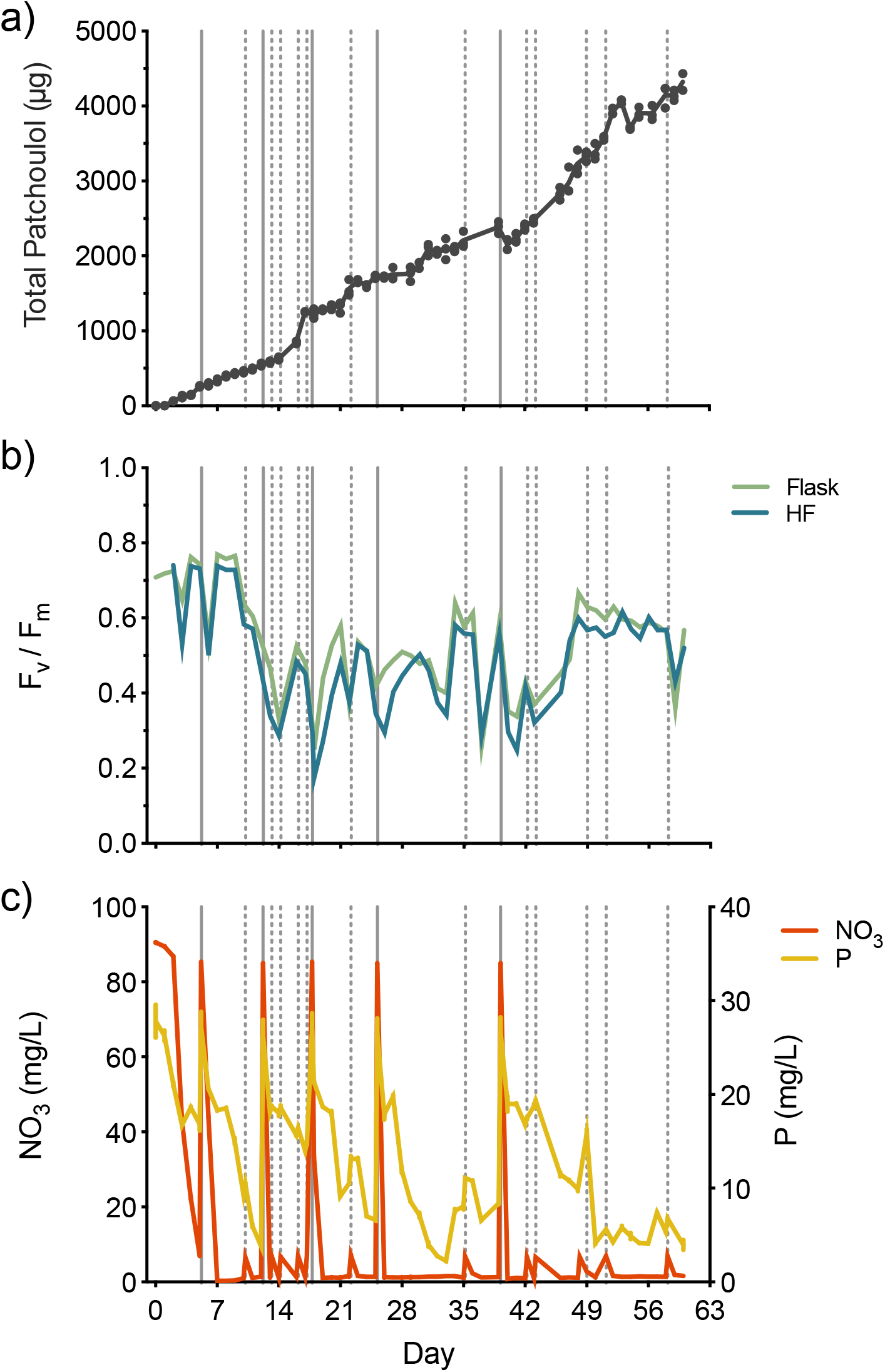
a) Total patchoulol produced by *C. reinhardtii* culture during 60-day cultivation coupled with the hollow-fiber cartridge. b) Maximum photochemical efficiency (*F*_v_/*F*_m_) of *C. reinhardtii* cells in the flask (green line) and the hollow fiber cartridge (blue) after 10 min of dark acclimation. c) Concentrations of nitrates (red) and total phosphorous (yellow) in the culture flask. Vertical dashed lines indicate when 100 mL of fresh TAP NO_3_ medium was added, while vertical grey solid lines show when 95% of the liquid culture (1.9L) was exchanged (solid grey lines) with fresh medium.

### Gas chromatography

To quantify the accumulation of patchoulol product in dodecane, 500 μL of solvent was sampled daily from the HF algal-culture setup. Solvent samples were analyzed using an Agilent 7890A gas chromatograph (GC) equipped with a DB-5MS column (Agilent J&W, USA) attached to a 5975C mass spectrometer (MS) with a triple-axis detector (Agilent Technologies, USA). A previously described GC oven temperature protocol was used ^26^. Patchoulol standard (18450, Cayman Chemical Company, USA) calibration curves in the range of 1–200 μM patchoulol in dodecane were used for quantification. 250 μM of α-humulene (item no. CRM40921, Sigma-Aldrich, USA) was added as an internal standard to each sample and patchoulol standard. All GC-MS measurements were performed in triplicate (n=3), and chromatograms were manually reviewed for quality control. Gas chromatograms were evaluated with MassHunter Workstation software version B.08.00 (Agilent Technologies, USA).

### Methanol toxicity assay

To determine whether residual amounts of methanol in dodecane inhibit the growth of the microalgae, a toxicity assay was performed. Briefly, 200μL of dense *C. reinhardtii* culture was added to 4.3 mL of TAP medium, and grown in 6-well microtiter plates with 500 μL solvent overlay consisting of dodecane with different amounts of methanol (0, 0.2, 0.5, 1, 2, and 5 ppm). Each methanol concentration was tested in biological triplicates (n=3), and the cultures were shaken at 120 rpm on a 12h:12h dark: light (approx. 150 μmol m^−2^ s^−1^) cycle for 7 d. After 7 d, the cell density of each culture was measured in triplicates (n=3) with an Invitrogen Attune NxT flow cytometer (Thermo Fisher Scientific, UK), as previously described^26^.

### Cross-flow nanofiltration for patchoulol concentration

Cross-flow filtration measurements were performed to determine the rejection values of patchoulol in methanol at 20 bar. A typical cross-flow nanofiltration rig was used for the cross-flow OSN experiments.^28^ The nanofiltration setup consisted of a feed tank, a pressure pump (Azura P 2.1L, Knauer, Germany), six stainless steel membrane cells, a recirculation gear pump (Micropump, INC), a 40-µm filter (Swagelok), a back pressure regulator (Swagelok), stainless steel connections (Swagelok), and FFKM grade sealing. The membrane cells were charged with three different membranes in duplicates, each having an effective area of 50.3 cm^2^. The membrane was a DuraMem 300 with an effective molecular weight cut-off of 300 g mol^−1^. Fig. 3c depicts the schematic diagram of the system. The total volume of the system was 220 mL, with a patchoulol concentration of 0.091 mg mL^−1^. Samples were collected from the permeate and the retentate streams, and then subjected to gas chromatography (GC-MS) to determine the patchoulol concentration. The rejection values were calculated according to Eqn 2.

**Figure 3.**
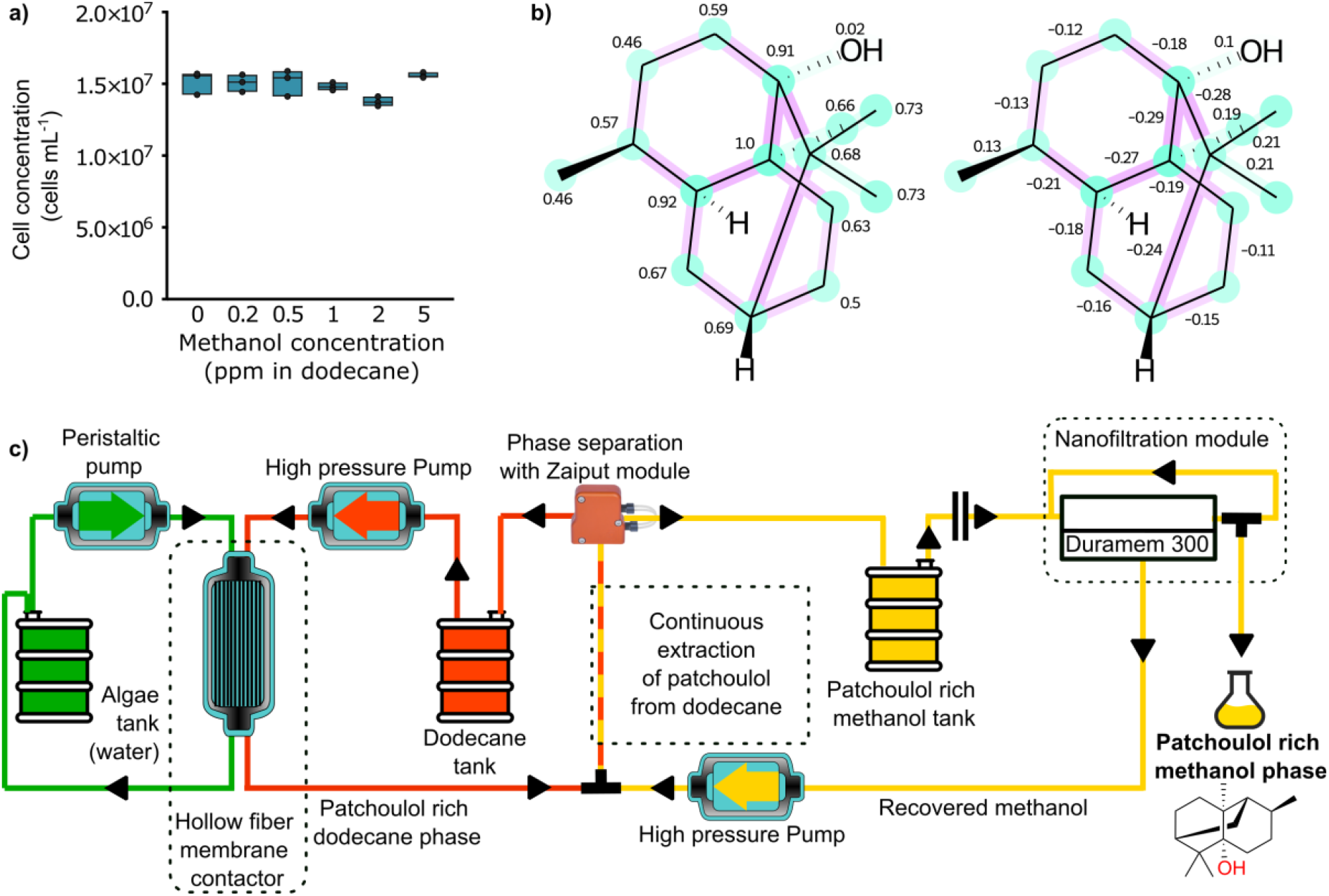
a) Cell concentration of *C. reinhardtii* at the end of the methanol toxicity assay: after 7 d cultivation of the algae with 10% (v/v) dodecane solvent overlay with different amounts of methanol contamination (n=3). b) Atomic and bond contributions for patchoulol nano-membrane rejection predicted by machine learning algorithm. Cyan color depicts positive contribution while purple color depicts negative contribution toward the final rejection value. c) schematic representation of the final process: the algae cultivation, dodecane extraction, methanol extraction, and nanofiltration modules. Arrows depict the direction of the flow of the liquid.

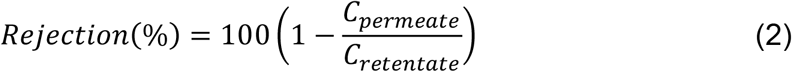

The machine learning model for the rejection prediction was adapted from literature^28, 29^. The model uses a message passing graph neural network to predict the rejection of the patchoulol and is trained on a large set of manually measured rejection values extracted from the OSN Database (www.osndatabase.com, accessed on 08.02.2022).^30^

### Nanofiltration

Nanofiltration experiments were performed to determine the rejection values of patchoulol in dodecane at 60 bar using different membranes. A typical dead-end filtration setup was used for OSN experiments^13^, consisting of a pressurized feed tank, membrane, nitrogen cylinder, and the necessary connections and sealings. The feed tank was stirred by a magnet to minimize the concentration polarization effect. Puramem S260, DuraMem 300 and GMT-oNF-2 membranes with an effective area of 7.4 cm^2^ were used. Samples were collected from the permeate and the retentate streams for GC-MS analysis to determine patchoulol concentrations.

### Partitioning coefficient measurement

The purpose of the partitioning coefficient 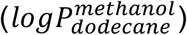 measurement was to determine the affinity of patchoulol between methanol and dodecane. The partitioning coefficient was determined by mixing the phases of methanol, dodecane, and patchoulol and allowing the system to reach a steady state. In a separation funnel, 5 mL methanol and dodecane containing approximately 4 ppm of patchoulol were first mixed and then phase-separated. Samples from the two phases were collected, and the patchoulol concentration was determined using GC-MS. The partitioning coefficient of the patchoulol was found to be 0.8 with respect to dodecane (Eqn 3).

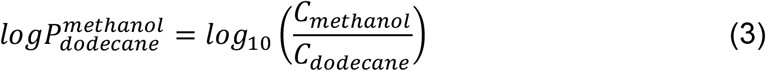

### Continuous membrane extraction measurements

Continuous extraction experiments were performed to determine the ability to remove patchoulol from dodecane using methanol by liquid-liquid extraction. The system consisted of flexible PFA tubings with a diameter of 1/16” (Perkin Elmer) and high pressure pumps (Vapourtech Ltd) controlled by a Vapourtech E-series unit. The mixer unit was a large diameter, static tubular module (Vapourtech), 20 mL in volume with an inner diameter of 3.2 mm. The phase separator unit was a Zaiput SEP-10 equipped with an IL-400 membrane. Separation of the organic and water phases were complete on the Zaiput up to 4 mL min^−1^ flow rate. The inlet stream consisting of patchoulol and dodecane (2 mL min^−1^) was mixed with another inlet stream of pure methanol (2 mL min^−1^). The unified stream was fed into the mixing unit and then into the Zaiput module, where phase separation occurred.

### Process Modeling

The purpose of modeling the extraction process was to obtain a comprehensive understanding of the efficiency and environmental impact of the entirety of the patchoulol extraction system. A dynamic MATLAB Simulink model was used to assess the Environmental (E) factor, solvent consumption, and economic sustainability of the coupled combination of bioprocess, extraction, and concentration using nanofiltration. The mathematical background for modeling the process is described in detail in the Supplementary section.

An important feature of the system is the slow enrichment of patchoulol from the microbial culture. The time required to generate a given amount of patchoulol is orders of magnitude higher than the time necessary to extract and concentrate the same amount from the solvent. Therefore, the generation and the extraction (up- and down-stream processes) were not directly continuously coupled in the model, and three different cases were examined:

#### Case 1: Semi-batch process with continuously coupled NF unit (Fig. S2)

The patchoulol in the dodecane was extracted with methanol when a specific concentration was reached. A given percentage of the patchoulol was extracted and concentrated continuously, then the extraction was halted, and the enrichment was resumed.

#### Case 2: Semi-batch process with detached NF unit

The same scenario as Case 1, but the solution in methanol was collected in a retentate loop or in a tank and separately concentrated to a given product concentration.

#### Case 3: Steady-state continuous operation

A sufficiently large volume of algae culture can maintain a constant concentration in the dodecane resulting in a fully continuous configuration. No dynamic simulation was required for cases 2 and 3.

Three green metrics, namely the E factor, solvent consumption, and economic sustainability, were used to assess the different configurations. The E factor is the ratio of the mass of the generated waste and the mass of the final product through the process (Eqn 4).

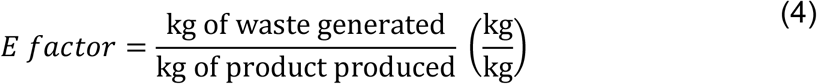

The waste generated in the described process was the consumed but not recycled solvent and the membrane. The consumption of the membrane was assumed to be 1 g year^−1^. The solvent consumption (SC) is the ratio of the mass of solvent waste generated and the mass of the isolated product (Eqn 5).

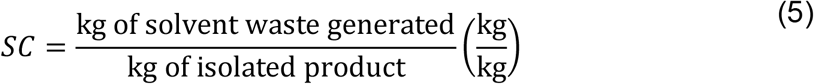

The economic sustainability (ES) is the ratio of mass of waste generated and the economic value of the isolated product (Eqn 6).

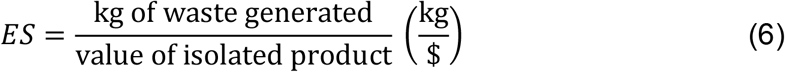

## Results and Discussion

### Hollow-fiber membrane contactor to extract algae-derived terpenoids

During the 60-day-long experimental period, a total of 4.3 mg patchoulol was extracted from a fluctuating 2 L reservoir of cultured engineered *C. reinhardtii* using a HF setup (Fig. 2a). In the first 42 days, extracted patchoulol averaged 398 ± 221 μg (mean ± SD) per week. In the subsequent two weeks, there was an increase in extraction amount, with 922 ± 26 μg patchoulol (mean ± SD) per week. Throughout the experimental period, the maximum photochemical efficiency (*F*_v_/*F*_m_) of the algal cells fluctuated considerably in the flask and HF cartridge, ranging from 0.25–0.77 and 0.18–0.74, respectively (Fig. 2b). The *F*_v_/*F*_m_ pattern of cells in the flask mirrored that of cells on the HFs, although the former generally had a slightly higher *F*_v_/*F*_m_ ratio, suggesting they were healthier. *F*_v_/*F*_m_ was highest in the first week, after which time it rapidly decreased and remained below 0.70; even after replacement of 95% of the medium. Overall, we found no evidence that patchoulol yield per day was significantly correlated with the *F*_v_/*F*_m_ (R^2^ = 0.009), indicating photosynthetic health was not related to sesquiterpenoid production.

Similarly, there was no evidence that patchoulol yield per day was significantly correlated with the concentrations of nitrate (R^2^ = 0.011) or total phosphorous (R^2^ = 0.014) in the culture media. Both nutrients gradually decreased in the first seven days of cultivation as the culture established (Fig. 2c). After this initial phase, nitrates were almost entirely depleted within 24 h following media changes, while total phosphorous concentration declined at a slower rate but did not reduce below 5 mg L^−1^.

After 4 weeks, we observed aggregation of algal cells on the HF strands. The settlement of cells meant that longer residence time at surface of the HFs. Patchoulol is partially water soluble^5^, and is found both in the cells and the culture medium^5^. Solvent contained within the HF should extract produced sesquiterpenoid from the culture medium as well as direct cell-solvent micelle interactions at the HF pores. It is possible that establishment of algal biofilm on the HF membrane enabled increased extraction efficiency into the solvent during the later phases of this trial.

Our goal was to investigate whether extraction of patchoulol from engineered algal culture was possible at all with such a HF cartridge setup. Optimization of culture conditions was outside the scope of the current study, and we relied on a relatively inefficient growth set-up using a stirred flask to simplify process testing. Future iterations could use cultivation systems optimized for maximal biomass productivity, such as small bubble chambers, wave bags, or membrane gas delivery^31^. The patchoulol yield (per liter of culture) obtained in the present study is similar to yields reported from solvent-culture two-phase cultivations in shake flasks; perhaps extraction efficiency would have continued to increase with increased biofilm establishment or with higher overall cell densities. Patchoulol titers of 70 μg/L culture after 5 days^32^ and ∼440 μg/L culture after 7 days ^5^ of cultivation in shake-flasks were previously reported in TAP medium. Here with the HF set-up, we were able to extract 470 μg patchoulol L^-1^ culture within 7 days (on days 42–49), however, significant optimizations could yet be implemented to increase yield rates of this process. Coupling an HF set-up to optimized algal cultivation units may enable higher-yield rates more amenable to industrial consideration. HF may also be used for direct gas exchange within cultures and increase efficiency of bio-reactor performance. Further parameter optimizations could also include increasing HF surface area to provide more contact area between the solvent and algae cells. In addition, further engineering to enhance yield of the cell catalyst will be crucial, as only a fraction of carbon (1 g L^− 1^ acetic acid) was converted to patchoulol in this experiment.

### Process development

The algal extraction process generated a dodecane phase with dilute concentrations of patchoulol. Further steps in downstream processing require the extraction of patchoulol from the dodecane phase. This separation is challenging because of similar chemical properties of both compounds. To reduce the amount of solvent waste generated in this separation, a continuous OSN system was implemented. A preliminary test indicated low patchoulol rejection, and low permeance of dodecane. The GMT-oNF-2 and the Puramem S260 membranes had a patchoulol rejection of 15%±10% and 1%±0.5%, respectively. The dodecane permeance value was 0.46 L m^−1^ h^−1^ bar^−1^ for the GMT-oNF-2 and 0.27 L m^−1^ h^−1^ bar^−1^ for the Puramem S260 membrane. The low permeance values are attributed to the high viscosity and large hydrodynamic volume of dodecane. The low rejection of patchoulol in dodecane can be explained by the similarity in their chemical structure. Both the patchoulol and the dodecane are nonpolar molecules, having high portioning 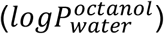 coefficients of 4.1 and 6.1, and contain mostly sp3 carbon atoms. The Van der Waals volumes of patchoulol and dodecane are also similar, with values of 236 Å^3^ and 216 Å^3^, respectively. These structural similarities imply having similar membrane–solute interactions resulting in the low rejection of patchoulol in dodecane, therefore, both molecules permeate through the membrane at a similar speed.^28^ Although replacing dodecane with another solvent with different chemical properties would be preferred for downstream applications, it is difficult to find other biocompatible solvents for product milking from living microorganisms.

We sought to select an immiscible solvent alternative, which would partition patchoulol from dodecane in a second-phase after extraction from the algal culture. Dodecane is immiscible with water, methanol, ethanol, acetonitrile, dimethyl sulfoxide, and dimethylformamide. Patchoulol has an approximately 10^4.1^ lower solubility in water than in dodecane and therefore water was ruled out. Dimethyl sulfoxide and dimethylformamide are not recommended solvents due to their toxicity, viscosity, and potentially damaging effect on the polymeric membranes.^33^ The remaining solvents, methanol and acetonitrile were, therefore, candidates for extraction.

We opted for methanol because it is a potentially sustainable solvent^34^, cheaper to produce, and it has a lower ICH limit (3,000 ppm) compared to acetonitrile. Moreover, recent studies related to the rejection prediction of small organic molecules in methanol allowed us to predict the rejection of patchoulol.^28, 29^ We also found that small amounts of methanol leaching into the dodecane phase would not perturb algal culture health (Fig. 3a). Using the open-access machine-learning prediction tool, the expected rejection of patchoulol in methanol is 100%. The measured rejection in our cross-flow OSN system confirmed the 100% rejection of patchoulol using Duramem 300 membrane. The results show that the recovery of methanol is possible with an OSN system, which can significantly reduce solvent loss and consequently the E factor of the process. Patchoulol loss during the OSN process is negligible due to its complete rejection from the membrane.

Fig. 3b shows the structure of patchoulol with the highlighted atom and bond-level contributions to the final rejection values. The expected nanofiltration membrane rejection is high because patchoulol is nonpolar and has many sp3 configuration carbon atoms, with relatively high positive atomic contributions and low negative bond contributions. Our machine learning-based prediction of 100% rejection matches the empirically observed rejection in methanol at 20 bar using Duramem membranes.

Fig. 3c shows a schematic diagram of the final process of the proposed patchoulol production system. First, the algae produce patchoulol and other high valued products using sunlight and wastewater. The process is continuously coupled with the dodecane extraction unit, which gradually extracts the oil-soluble components from the algae. Once the patchoulol concentration in the dodecane is high enough, the methanol extraction from the dodecane can start. The extracted patchoulol and methanol are collected and then filtered through an OSN system at 20 bar using a Duramem 300 membrane. The rejection of the patchoulol in this system is 100%; therefore, no loss of patchoulol to the permeate stream occurs. The concentrated patchoulol-methanol solution (>10 mg mL^−1^) is collected at the retentate side of the membrane module.

### Analysis of nanofiltration-coupled configurations and their sustainability

The main engineering parameters examined in sustainability Case 1 modeled here are the flow rate of the methanol, the area of the membrane, and the applied pressure. The initial concentration of patchoulol in the dodecane was set to 11 μg mL^−1^ (the maximum concentration examined in Fig. 2A). First, we investigated the effect of the methanol flow rate on the system and the output. The dodecane flow was set to 2 mL min^−1^. Owing to the theoretical maximum capacity of the Zaiput module (10 mL min^−1^), we selected two methanol flow rates to be examined: 2 mL min^−1^ (Case 1A) and 8 mL min^−1^ (Case 1B). The concentration profiles in the dodecane, the methanol inlet stream (NF feed) and in the NF efflux (retentate) are shown in Fig. 4.

**Figure 4.**
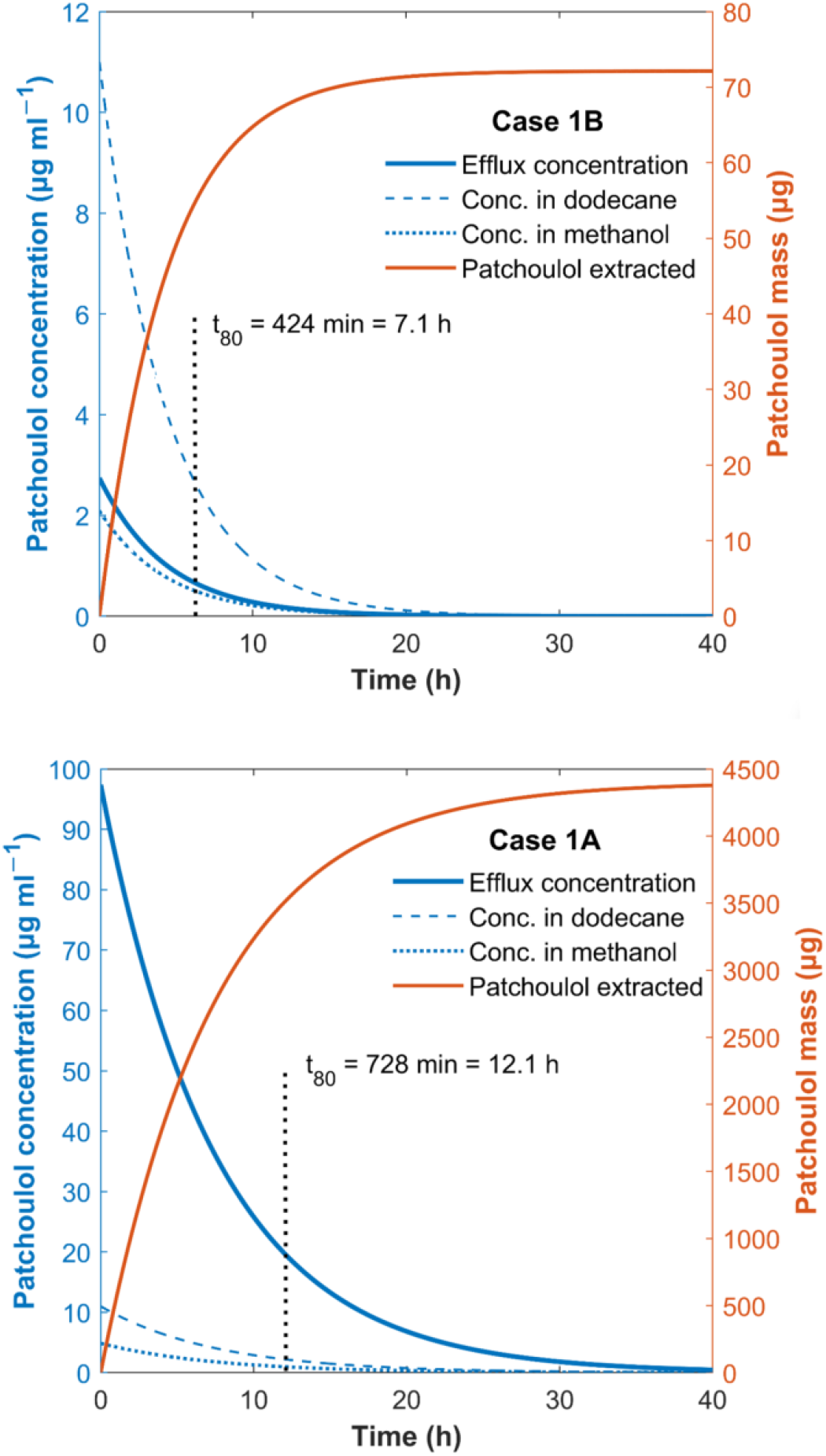
Patchoulol concentrations in dodecane and methanol over time for a) Case 1A and b) Case 1B with methanol flow rates of 2 mL min^−1^ and 8 mL min^−1^, respectively.

Lower methanol flow rate resulted in higher output patchoulol concentration at the same operating pressure (Fig. 4) because the solution leaving the Zaiput module was less diluted. Examining the time required to extract 80% of the patchoulol from the dodecane solution: case 1A requires 728 min while case 1B 424 min. However, the enrichment period to yield patchoulol from the microbial culture from 2.2 μg mL^−1^ to 11 μg mL^−1^ in dodecane was found to be 62.9 d in our HF-culture setup, so the difference in extraction time is negligible compared to the total required time to accumulate product. The reduced extraction time cannot make up for the deterioration of environmental factors caused by the dilution of the efflux solution. Increasing the methanol flow rate results in an excessively diluted solution, and the decreased extraction time has no significant effect on productivity as the enrichment phase from the microbial culture is by far the slowest process.

Using sensitivity analysis, the effects of various engineering and system parameters on the green metrics of the process were mapped out. We examined case 1A, varying the extraction rate into methanol and the initial patchoulol concentration in dodecane on levels 20%, 40%, 60%, 80%, and 2 μg mL^−1^, 6 μg mL^−1^, 11 μg mL^−1^, respectively. We calculated economic (E) factor (comprised of both solvent and membrane waste) and economic sustainability for the different cases, with their respective results shown in Fig. 5a and b.

**Figure 5.**
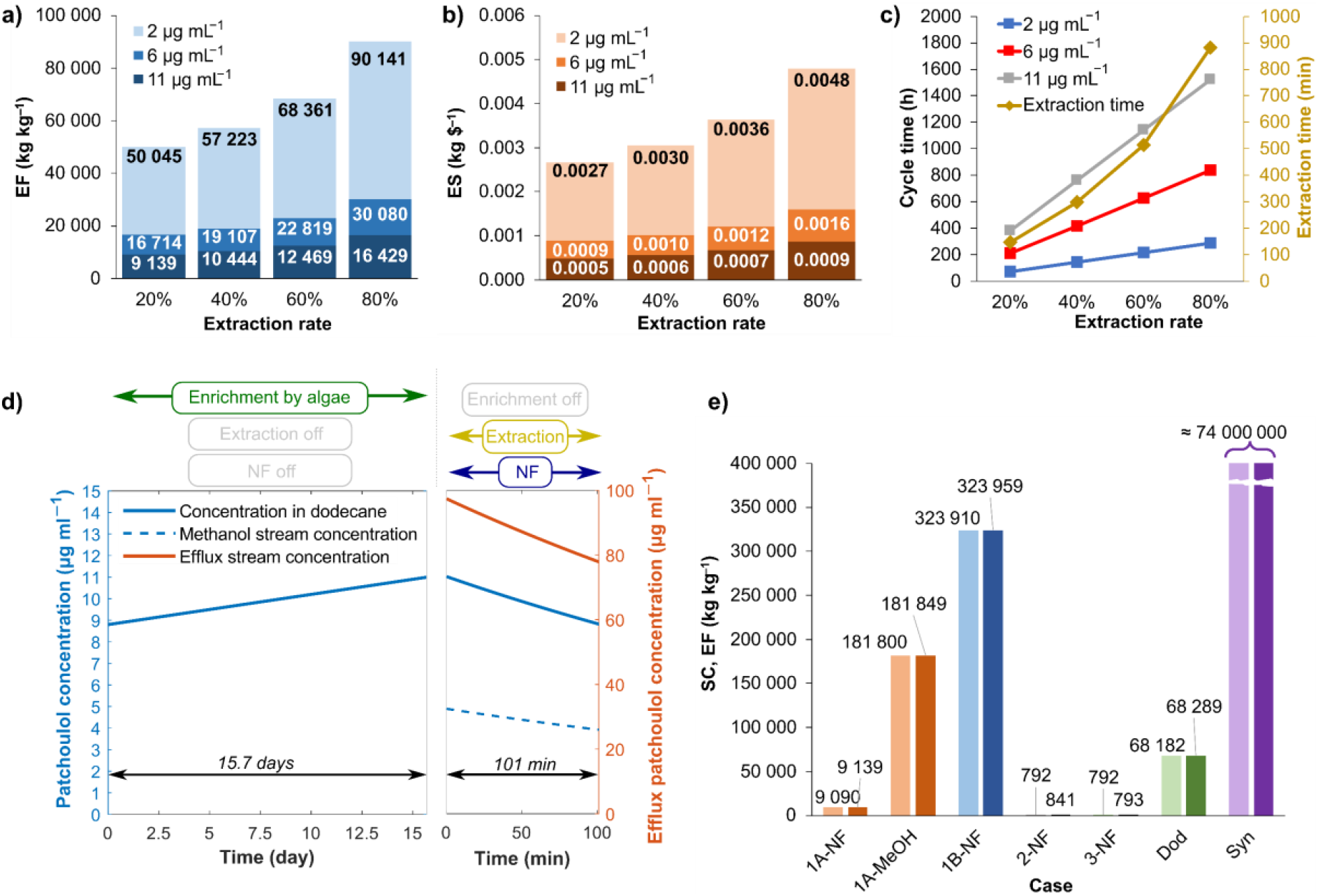
a) Sensitivity analysis of the E factor in case 1A. b) Sensitivity analysis of the economic sustainability, (c) cycle time and extraction time, (d) process cycle for case 1A, showing steps of the process and the timeframe for each operation. e) Comparison of green metrics for main cases of process configuration.

Not surprisingly, we found that lower extraction rates of patchoulol from dodecane into methanol and higher initial patchoulol concentrations in dodecane improved the sustainability features of the process. As extractions are continued for longer time frames (higher extraction rate), the methanol solution flow becomes more diluted, leading to higher solvent consumption (waste), therefore lower extraction rates are advantageous for process efficiency. Fig. 5c illustrates that the time cycle describing the semi-batch operation increases both with extraction rate and initial concentration values due to the longer enrichment times. However, this does not result in a less efficient process since the average molar efflux of patchoulol is almost the same in all cases, the rate-limiting step is always the patchoulol enrichment phase into dodecane from the microbial culture. The rate of patchoulol generation is orders of magnitude lower than any of the examined extraction procedures, which is what determines the overall productivity of the process. Since the rate of generation does not change with time, longer time cycles do not decrease the amount of patchoulol that can be extracted in a timeframe.

In a real-life situation, the parameters examined above have a significant effect on the cost of the process. Therefore, economic optimization is necessary to find industrial relevance. Based on the results illustrated in Fig. 5a, we chose 11 μg patchoulol mL^− 1^ dodecane as the semi-optimal initial concentration value and 20% extraction rate to compare Case 1 with other cases. Fig. 5d shows the modeled concentration values for a process cycle corresponding to the chosen system parameters. The very first process cycle is preceded by a run-up period of patchoulol enrichment to achieve the 8.8 μg mL^−1^ minimal concentration. The run-up time for the case presented here is 62.9 days.

### Comparative assessment of different configurations

In **case 1** (with the continuously coupled NF unit), it is challenging to achieve high efflux concentrations from the dilute methanol solution. Suppl Fig. 3 demonstrates the output concentration as a function of pressure for four different membrane areas. The pressure interval where high concentrations can be reached is very narrow, and operating at such values would result in losing the robustness of the process due to high sensitivity. The solubility of patchoulol in ethanol is 16 mg mL^−1^ 35; assuming a similar solubility in methanol (with a slight decrease due to the higher polarity of methanol), we can theoretically produce solutions of around 10 mg mL^−1^; however, the configuration described in case 1 does not allow that. A very precise pressure value must be maintained, which is not practically possible, although a new configuration with a retentate cycle and a buffer tank (case 2) can solve this problem. This modification improves the efficiency, E factor, and solvent consumption of the system. In **case 2**, the nanofiltration unit is decoupled from the extraction system. The methanol solution of patchoulol can be collected in a container or in a retentate loop and can be concentrated in a separate unit operation, independently from the previous extraction steps. In this case, we do not have to specify the membrane area and the pressure, as these are usually determined from economic metrics. In the comparative assessment of sustainability features, we assume that the methanol solution is concentrated to 1 mg mL^−1^.

**Case 3** describes a completely continuous configuration. Here, we assume that a sufficiently large algae culture can maintain a constant concentration of 11 μg patchoulol mL^−1^ in the dodecane solution while operating the methanol extraction cycle parallelly. Similar to case 2, we assume that a retentate cycle or a buffer tank enables robust concentration of the mixture to 1 mg mL^−1^ with an adequate nanofiltration unit. Because of this, the green metrics of cases 2 and 3 will be similar, with the slight difference that the higher molar efflux of case 3 enables more efficient membrane usage within its life cycle, thereby reducing the environmental factor.

The comparison of solvent consumption and E factor for different system configurations and cases is illustrated in Fig. 5e.^36^ Cases *1A-NF* and *1B-NF* represent the entire process with coupled NF for 2 mL min^−1^ and 8 mL min^−1^ methanol flow rates, respectively. *1A-MeOH* represents case 1A without the nanofiltration unit and methanol recycling (as if we consider the initial methanol solution the product), while *2-NF* and *3-NF* stand for cases 2 and 3. For a comprehensive comparison, we also considered the case where the dodecane solution is the final product (*Dod*), and calculated the green metrics for a synthetic route described in literature.^37^

Fig. 5e shows that raising the methanol flow rate drastically decreases the sustainability features due to the highly diluted efflux. Also, the nanofiltration step, in coupled configuration, greatly decreases the SC (solvent consumption) and ES (economic sustainability) values, but it is still not as efficient as cases 2 or 3, where the concentration of the final product is arbitrarily high. The only difference between cases 2 and 3 is the slightly lower EF-value of the latter, showing a more efficient membrane life cycle exploitation for the scaled-up fully continuous configuration. In a pilot plant or on an industrial scale, the efficient use of membrane modules can be crucial. The implementation of a completely continuous operation of the bioprocess can not only increase sustainability but can also enhance the robustness of the system and process control by eliminating the time factor. This approach, however, requires significantly larger algae cultivation set-ups or algae strains with increased productivity.

## Conclusions

In this work, we showed that a system comprised of a HF membrane contactor, membrane-based liquid-liquid extraction, and OSN can be used for efficient in-line extraction and concentration of patchoulol from engineered algal culture. The patchoulol titers obtained in the dodecane reservoir with the HF setup were similar to those previously reported for traditional two-phase extractions in flasks, however, at much longer timescales. While the obtained patchoulol yields were reasonably high, additional modifications to the HF setup, such as using a larger-surface-area HF cartridge and coupling this to an optimized algal cultivation unit, are likely to increase yields even further. Further developments in strain engineering to increase yields will also likely improve titers. This process would function well with other engineered microbes, which could also boost process efficiencies. We also showed the implementation of “separation by design” green engineering principle nanofiltration. Molecular modeling and empirical testing indicated expanding extraction with methanol could enable efficient, low energy extraction of the target product, and we present the efficiency of such processes for many modeled cases. Our proposed system presents a low-waste and low-energy means to enrich patchoulol extracted from the microbial culture. We demonstrated that the decoupled configuration and operation of the membrane concentration unit helps to reduce the environmental footprint of the system, maximizes efficiency, and enables an inherently robust downstream process. Further implementations of this concept will require improvements in algal-HF membrane contact and target metabolite productivity to maintain a fully continuous process.

## Author Contributions

Sebastian Overmans: Conceptualization, Microbial cultivation, HF membrane implementation, Gas Chromatography, Data analysis, Visualization, Writing – original draft

Gergo Ignacz: Conceptualization, Visualization, Data curation, Formal analysis, Machine learning, Nanofiltration, Methodology, Writing – original draft

Aron K. Beke: Process modeling, Data analysis, Visualization, Writing – original draft

Jiajie Xu: Conceptualization, Material sourcing, Writing – review & editing

Pascal Saikaly: Conceptualization, Writing – review & editing, Funding Acquisition, Project Administration

Gyorgy Szekely: Conceptualization, Resources, Methodology, Investigation, Writing – review & editing, Supervision, Funding acquisition, Project administration.

Kyle J. Lauersen: Conceptualization, Funding Acquisition, Project Administration, Resources, Methodology, Investigation, Writing – original draft

## Conflicts of interest

There are no conflicts to declare.

## Acknowledgements

We would like to express special thanks to Najeh Kharbatia of the KAUST Analytical Core Labs for helpful early discussions, Chandrasekaren Lakshmipathy, Abdulkhalik Khalifa, and Abdullah Alabdullatif of KAUST Lab Equipment Maintenance (LEM) team for assistance in upgrading and initializing the GC-FID-MS unit. The authors acknowledge Prof. Dr. Ralph Bock for providing *C. reinhardtii* UVM4, obtained under material transfer agreement between KAUST and the Max-Planck-Institut für Molekulare Pflanzenphysiologie, Potsdam. The research reported in this publication was supported by the KAUST Impact Acceleration Funds program (grant 4238), and KAUST baseline funding awarded to Kyle Lauersen and Gyorgy Szekely. Figure 1 was produced by Ana Bigio, scientific illustrator.

## Notes

### Competing Interest Statement

The authors have declared no competing interest.

