## Supplementary Information for "Continuous extraction and concentration of secreted metabolites from engineered microbes using membrane technology"

Electronic Supplementary Information for

### Contents of this file

Suppl. Figures 1–3

Suppl. Methods: Process Modeling

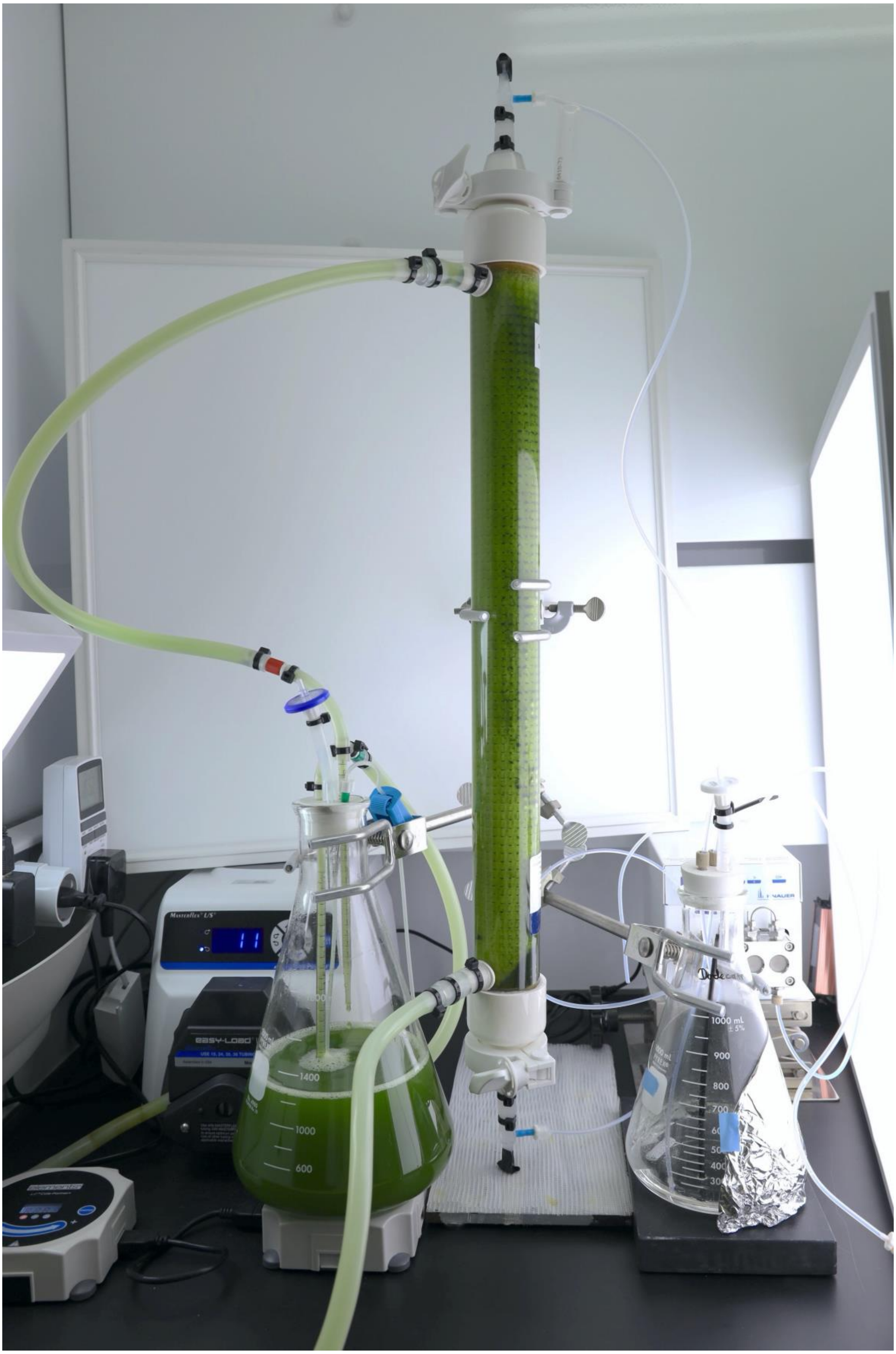


**Suppl. Figure 1.** Photo of hollow-fiber setup used to extract patchoulol from the alga *C. reinhardtii.*

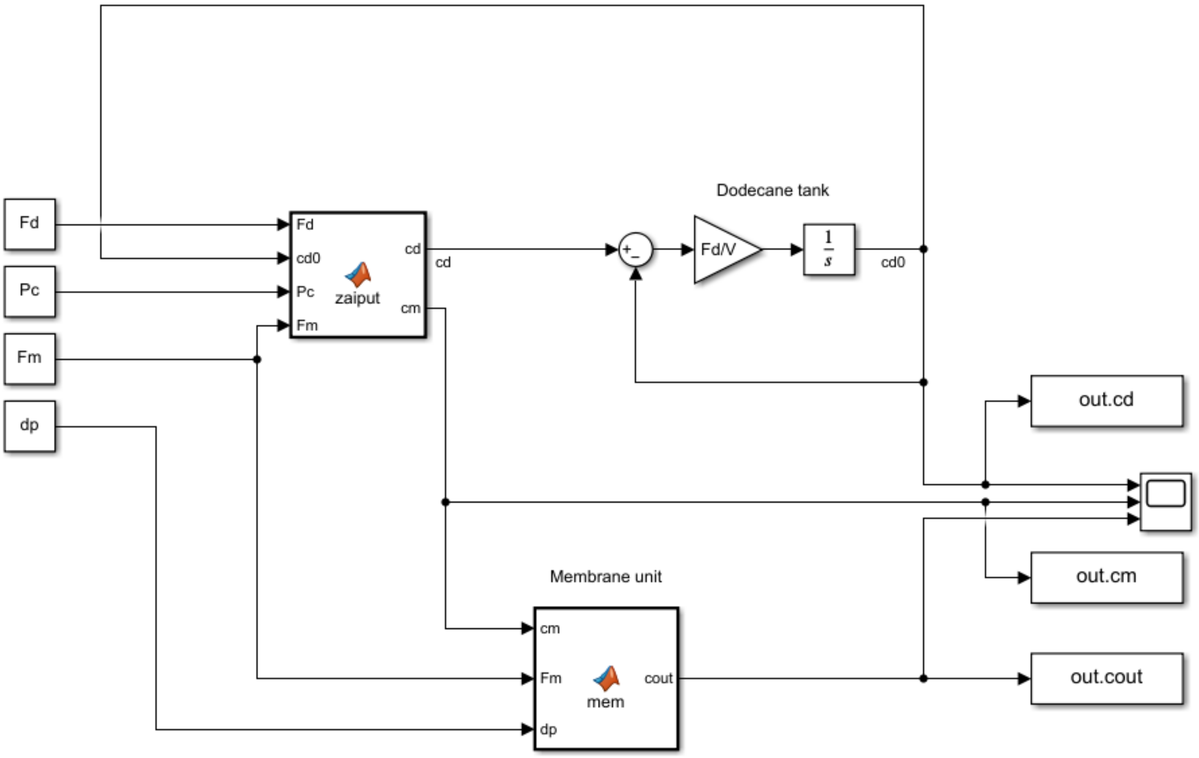


**Suppl. Figure 2.** MATLAB Simulink model of case 1 of process modeling.

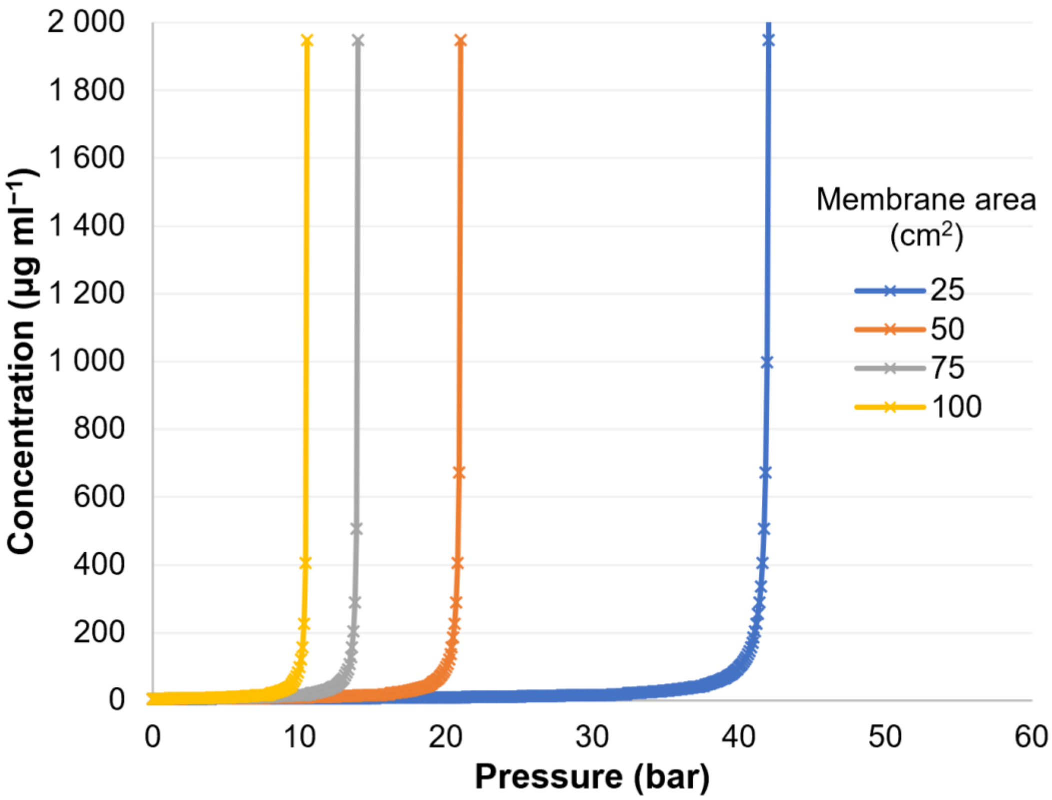


**Suppl. Figure 3.** Initial efflux concentration as a function of pressure difference. Only a narrow pressure interval provides sufficient output; however, this is not a robust solution from an engineering point of view.

**Suppl. Methods: Process Modeling**

A constant mass species generation term describes the extraction of patchoulol from the aqueous medium into dodecane. The dodecane was circulated in a closed loop with a constant volume of 400 mL. Therefore, the patchoulol concentration increased linearly in time without extraction (see Fig. 2a).

The experimental values were obtained for different batches of algae. Therefore, the temporal concentration gradients changed over time. To model the system, an arbitrary value from the measured range was chosen for the generation of the product:

$$\frac{\partial c_d}{\partial t} = 0.140 \frac{\mu g}{mL \cdot day} \tag{1}$$

The dodecane tank was modeled by a continuous stirred tank-reactor (CSTR) behavior, where the bulk concentration equals the tank output concentration ($c_d^0$). The species mass balance equation describing the dynamic behavior of the tank is:

$$\frac{dc_d^0}{dt} = \frac{F_d}{V}(c_d - c_d^0) \tag{2}$$

, where $c_d$ is the concentration of the patchoulol in the dodecane stream leaving the Zaiput, $F_d$ is the flow rate of the dodecane solution, and $V$ is the volume of the dodecane in the tank ($V = 400\,mL$).

The dodecane-methanol extraction was modeled by assuming equilibrium concentrations in the output dodecane and methanol streams, determined by the corresponding $logP_{dodecane}^{methanol}$ value. The solubility of methanol in dodecane was determined to be 0.2 w% and not to be toxic to the algae (see Fig. 3a). Therefore, removing methanol from dodecane after the extraction step was not necessary. Eqn 3, $c_m$ is the concentration in the methanol solution from the Zaiput to the NF, $F_m$ is the flow rate of the methanol solution, $P_c$ is the partition coefficient derived from the measured $logP_{dodecane}^{methanol}$.

$$c_m = \frac{F_d c_d^0}{F_d P_c + F_m} \tag{3}$$

The nanofiltration module (NF) was implemented to concentrate the initial dilute solution of methanol. To simplify the modeling of the membrane module, the pressure-dependence of the flux was assumed to be linear, determined by the permeance of the membrane ($P$), as implemented in Eqn 4. As the rejection of patchoulol is 100%, the permeate of the NF unit is pure methanol. The concentration of the NF efflux (retentate) leaving the nanofiltration unit:

$$c_r = \frac{F_m c_m}{F_m - PA\Delta p} \tag{4}$$

In Eqn 4, $A$ is the membrane area, $\Delta p$ is the pressure difference. The dodecane flow was 2 mL min$^{-1}$ in all cases of modeling, the membrane was modeled as a Duramem 300 unit with the following permeance:

$$P = 1.9 \times 10^{-3} \frac{mL}{cm^2 \cdot min \cdot bar} \tag{5}$$

MATLAB Simulink was used to create a comprehensive model of the extraction and nanofiltration processes (Suppl. Fig. 2).
